# Genetic epidemiology of variants associated with immune escape from global SARS-CoV-2 genomes

**DOI:** 10.1101/2020.12.24.424332

**Authors:** Bani Jolly, Mercy Rophina, Afra Shamnath, Mohamed Imran, Rahul C. Bhoyar, Mohit Kumar Divakar, Pallavali Roja Rani, Gyan Ranjan, Paras Sehgal, Pulala Chandrasekhar, S. Afsar, J. Vijaya Lakshmi, A. Surekha, Sridhar Sivasubbu, Vinod Scaria

**Author notes:** Authors contributed equally and would like to be known as joint first authors (Bani Jolly, Mercy Rophina). Address for correspondence: Vinod Scaria.

## Abstract

Many antibody and immune escape variants in SARS-CoV-2 are now documented in literature. The availability of SARS-CoV-2 genome sequences enabled us to investigate the occurrence and genetic epidemiology of the variants globally. Our analysis suggests that a number of genetic variants associated with immune escape have emerged in global populations.

## Text

Antibodies are one of the emerging therapeutic approaches being explored in COVID-19. These antibodies typically target the receptor-binding motif or structural domains of the Spike protein of SARS-CoV-2, in an attempt to inhibit binding of Spike protein with the host receptors. Cocktails of antibodies which target distinct structural and functional domains of spike proteins are also being currently developed considering redundant mechanisms of targeting the virus and therefore minimising escape mechanisms. Genomic documentation of the spread of SARS-CoV-2 across the globe has provided unique insights into the genetic variability and variants of functional consequence. In-depth studies in recent months have unravelled a wealth of information on the immune response in COVID-19 and offered insights into the development of therapeutics.

Recent investigations suggest a number of genetic variants in SARS-CoV-2 are associated with immune escape and/or resistance to antibodies. Their structural and functional features and mechanisms of immune evasion are also being extensively studied (*1*). The natural occurrence and genetic epidemiology of these variants across the global populations are poorly understood. We were motivated by the wide availability of SARS-CoV-2 genomes from across the world and the increasing numbers of genetic variants suggested to contribute to escape from antibody inhibition.

We analysed a comprehensive compendium of genetic variants associated with immune escape and curated by our group from literature and preprint servers (*2*). This compendium included 120 unique variants reported in literature. To understand the genetic epidemiology of these variants in the global compendium of genomes, we compiled the dataset of 265,079 SARS-CoV-2 from GISAID (as of 17 December 2020) (*3*) apart from 1,154 genomes sequenced in-house (BioProject ID: PRJNA655577). Genome sequences with more than 5% Ns, more than 10 ambiguous nucleotides, higher than expected divergence and mutation clusters were excluded from the analysis. After quality control, the final dataset encompassed 240,133 genomes from 133 countries. Only countries with at least 100 good quality genome submissions were considered for the analysis.

86 of the 120 genetic variants associated with immune escapes were found in a total of 26,917 genomes from 63 countries (**Figure 1A**), out of which 9 variants had >1% frequency in the respective countries. Phylogenetic analysis was performed following the Nextstrain protocol for a total of 3,679 genomes, including 1,501 randomly selected genomes having these variants (**Figure 1B**) (*4*). Homoplasies were identified in the phylogeny using HomoplasyFinder (*5*). Out of 86, 43 variant sites were found to be homoplasic, suggesting they could emerge independently in different genetic lineages, out of which 9 were found to be at >1% frequency in at least one of the countries analysed.

**Figure 1.**
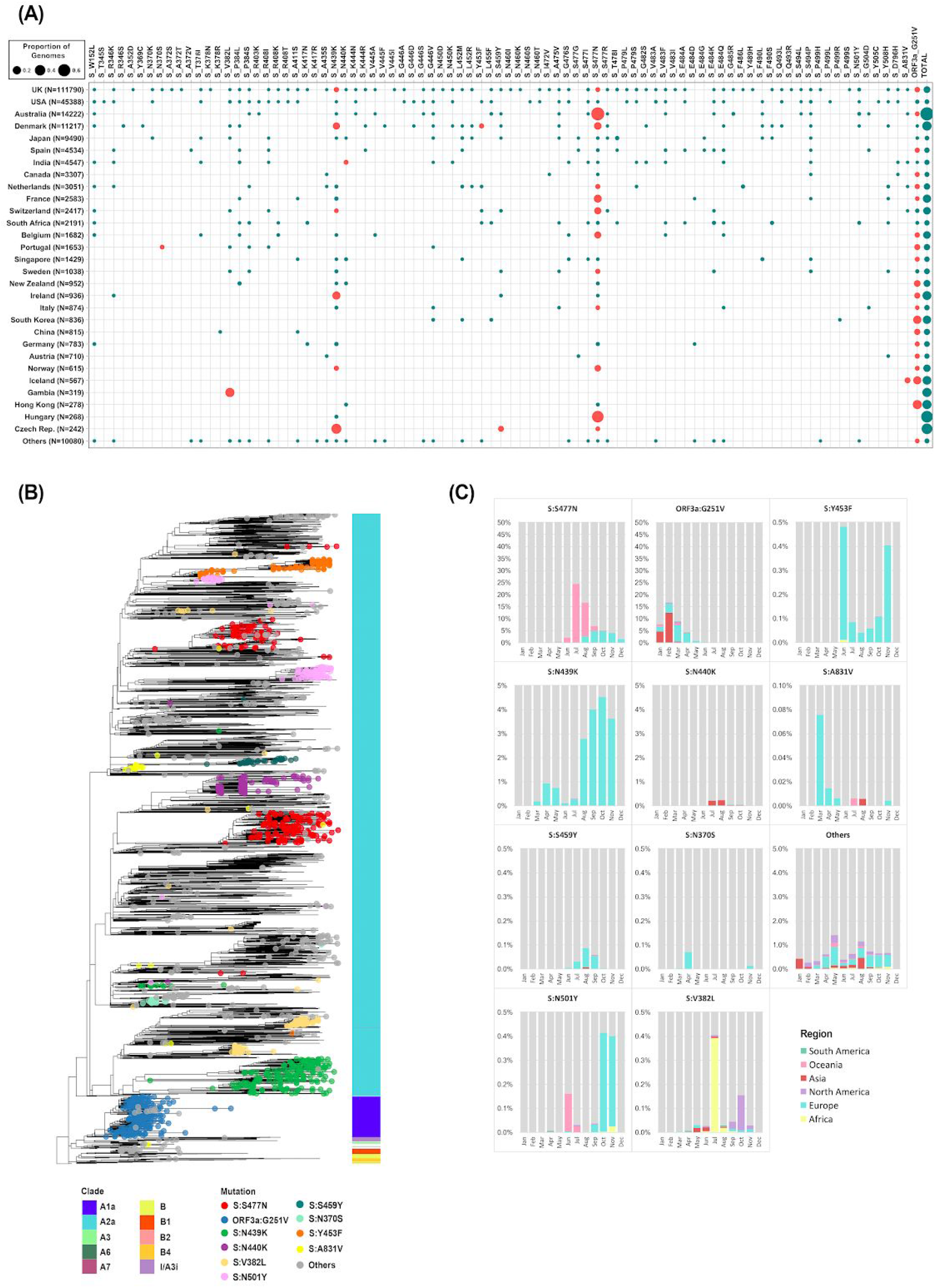
(A) Variant frequencies of the immune escape variants in genomes of SARS-CoV-2. The total number of genomes analyzed from each country is specified. Variants with frequency >1% in the respective countries are highlighted in red. (B) Global phylogenetic context of the variants. The vertical bar indicates the clade assigned according to the Nextstrain nomenclature (C) Time-series data on prevalence for the genetic variants showing the region-wise proportion of genomes per month for the variants

Out of 14,222 genomes analysed from Australia, 24 immune escape associated variants mapped to 9,895 genomes (70%). Of significant frequency was the S:S477N variant which was present in 9,541 genomes (67%) from Australia. High frequency of this variant was also found in a number of other countries particularly in Europe. S:N439K was also found at high frequencies in genomes from a number of countries in Europe (*6*).

S:N501Y, one of the variants in the recently reported emergent SARS-CoV-2 lineage from the United Kingdom, was present in a total of 290 genomes, including genomes from the United Kingdom, Australia, South Africa, USA, Denmark and Brazil (*7, 8*). All 7 genomes from South Africa having S:N501Y also had the S:E484K variant and S:K417N was present in 2 of these genomes (*9*).

The ORF3a:G251V variant was also found to be prevalent across global genomes, with the highest frequencies in Hong Kong and South Korea. This variant is also one of the defining variants for the Nextstrain clade A1a (GISAID Clade V) (**Figure 1B**).

19 of the 86 genetic variants were found in genomes from India (**Supplementary Figure**). The S:N440K variant was found to have a frequency of 2.1% in India and a high prevalence in the state of Andhra Pradesh (33.8% of 272 genomes). The variant site was homplasic and the variant was found in genomes belonging to different clades and haplotypes. Time-scale analysis suggested the variant emerged in recent months (**Figure 1C**). The S:N440K variant was also reported in a case of COVID-19 reinfection from North India (*10*).

Put together, our analysis suggests that a number of genetic variants which are associated with immune escape have emerged in global populations, some of them have been found to be polymorphic in many global datasets and a subset of variants have emerged to be highly frequent in some countries. Homoplasy of the variant sites suggests that there could be a potential selective advantage to these variants. Further data and analysis would be needed to investigate the potential impact of such variants on the efficacy of different vaccines in these regions.

## Acknowledgements

Authors acknowledge Disha Sharma and Abhinav Jain for the analysis of in-house genomes and the researchers, originating and submitting laboratories of the sequences retrieved from GISAID (https://doi.org/10.6084/m9.figshare.13365503.v2). BJ and MKD acknowledge a research fellowship from the Council of Scientific and Industrial Research (CSIR India). The funders had no role in the study design or the decision to publish.

---

**Supplementary Figure.**
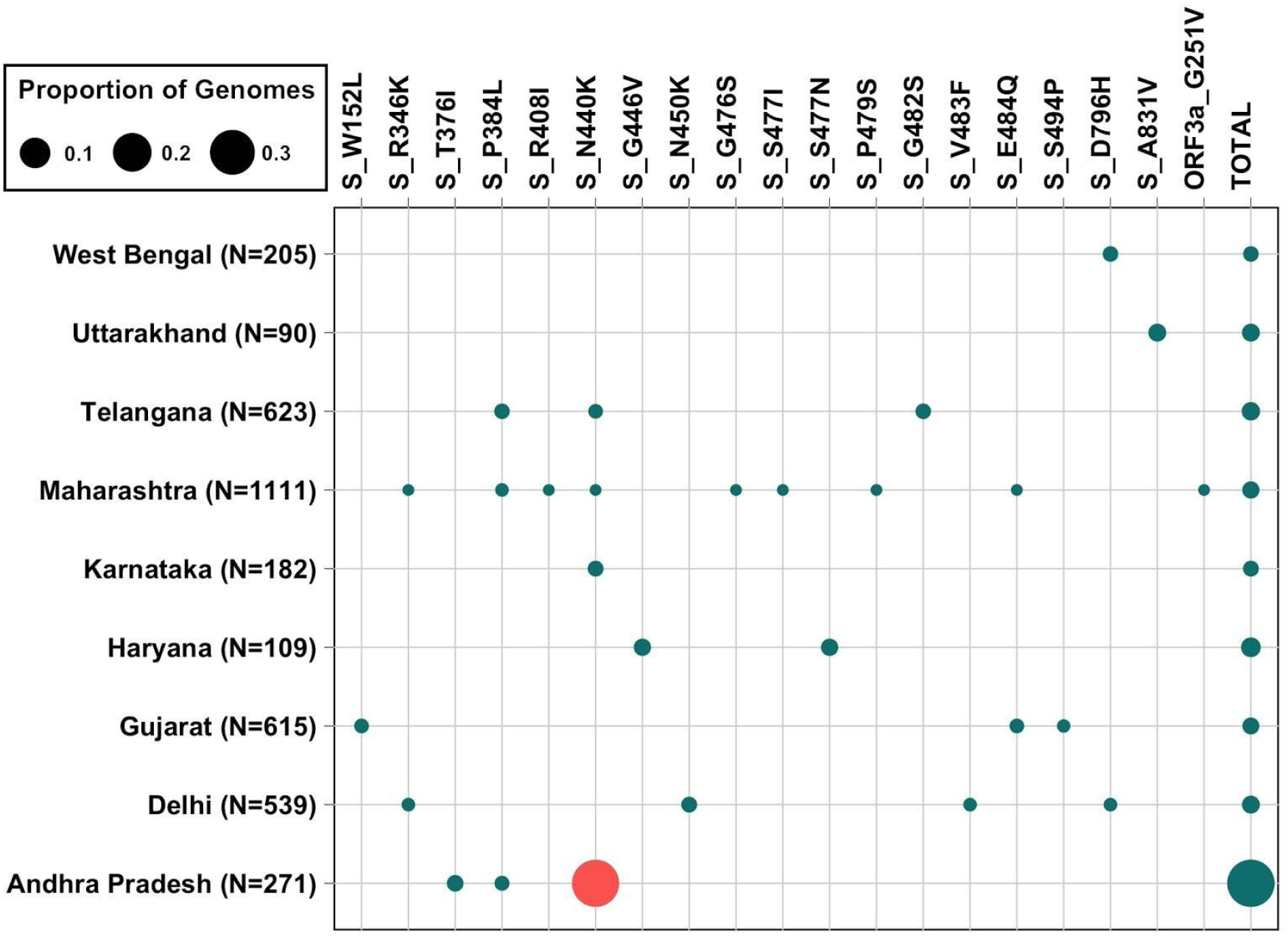
Variant frequencies of the immune escape variants in genomes isolated from different states in India.

